# A pan-cancer landscape of interactions between solid tumors and infiltrating immune cell populations

**DOI:** 10.1101/192286

**Authors:** David Tamborero, Carlota Rubio-Perez, Ferran Muiños, Sabarinathan Radhakrishnan, Josep M Piulats, Aura Muntasell, Rodrigo Dienstmann, Nuria Lopez-Bigas, Abel Gonzalez-Perez

**Author notes:** These authors contributed equally to this work (David Tamborero, Carlota Rubio-Perez).

## Abstract

Throughout their development, tumors are challenged by the immune system and acquire features to evade its surveillance. A systematic view of these traits is still lacking. Here, we identify genomic and transcriptomic traits associated to the immune-phenotype of 9,403 tumors of 29 solid cancers. In highly cytotoxic immune-phenotypes we found tumors with low clonal heterogeneity enriched by alterations of genes involved in epigenetic regulation, ubiquitin mediated proteolysis, antigen-presentation and cell-cell communication, which may drive resistance. Tumors with immune-phenotypes with mid cytotoxicity present an over-activation of processes involved in invasion and remodeling of neighboring tissues that may foster the recruitment of immune-suppressive cells. Tumors with poor cytotoxic immune-phenotype tend to be of more advanced stages and present frequent alterations in cell cycle, hedgehog, beta-catenin and TGF-beta pathways, which may drive the immune depletion. These results may be exploited to develop novel combinatorial targeting strategies involving immunotherapies.

---

Solid tumors and the immune cells infiltrating them interact in a dynamic equilibrium that shapes the progression of the disease. Indeed, tumor evasion of the immune surveillance is a hallmark shared by all types of cancer^1^. In this process, the features that protect cancer cells from the action of a cytotoxic immune infiltrate or promote the suppression of this infiltrate are positively selected. Although several mechanisms of immune evasion have been experimentally evinced for some tumor types^2–4^, we still lack a comprehensive view of routes of tumor immune escape. Some of these mechanisms –such as the activation of immune checkpoints– have been therapeutically exploited, with a remarkable clinical success across a variety of cancer types^4^. Therefore, the discovery of features acquired by tumors in response to the immune cells that surround them may open up new strategies to treat the disease.

The genomic and transcriptomic sequencing of large cohorts of tumors generated by international projects —such as The Cancer Genome Atlas (TCGA)– have provided the opportunity to map the molecular traits of cancer in unprecedented detail. Furthermore, the abundance of immune cells in the infiltrate of tumors can be estimated via bioinformatics approaches that exploit the fact that tumor samples are admixtures of cancer cells and their microenvironment^5^. Although several recent studies using these approaches have explored whether molecular features of the tumors are associated with specific characteristics of the immune infiltrate^6–10^, a comprehensive pan-cancer landscape of the interactions between the tumor and the immune cells is still lacking.

We consider the immune infiltration pattern of tumors –ignoring differences of immune competence between individuals– to be the result of three components, namely the tissue of origin, the cancer type and the specific molecular features of the tumor. Here, we evaluated the contribution of each of these sources to the infiltration pattern –represented by a comprehensive set of immune cell populations– of solid tumors. Our results show that the heterogeneity of the immune infiltrate cannot be explained solely by the two first components. Therefore, we set out to study –to the best of our knowledge– the broadest array of tumor intrinsic features (clinical, genomic and transcriptomic) that are related to their capabilities to evade the immune system in a pan-cancer cohort of 9,403 solid tumors. As a result, we delineate different scenarios of quantity and quality of the immune infiltrate of these malignancies and identified the landscape of oncogenic events associated to them. We reveal that tumors develop a very distinct set of molecular characteristics when progressing in these diverse immune scenarios, and provide novel insights of the interactions between tumors and immune cells in solid cancers.

## RESULTS

Figure 1 summarizes the analyses described in this paper. We define the immune infiltration pattern of each tumor as the relative abundance of an array of cell populations of the adaptive and innate immune system. After rigorous evaluation the performance of several tools designed to compute this abundance from the RNA-seq data of tumor samples (Supp. Methods), we selected a gene set enrichment analysis method (GSVA)^11^. To represent the immune infiltration pattern, we chose the most specific immune cell populations whose computed abundance across tumors produced biologically sound results given the gene sets available from several sources (Suppl. Methods). These were B cells, eosinophils, macrophages, mast cells, NK CD56bright cells (NKbright), NK CD56dim cells (NKdim), neutrophils, T helper cells (Th), central memory T cells (Tcm), effector memory T cells (Tem), follicular helper T cells (Tfh), activated dendritic cells (aDC), immature dendritic cells (iDC), activated CD8 T cell (CD8+ T), gamma delta T cells (Tgd) and regulatory T cells (Treg). The rationale for the selection of the gene sets employed for each of them (Table S1) is also discussed in the Supp. Methods.

**Figure 1.**
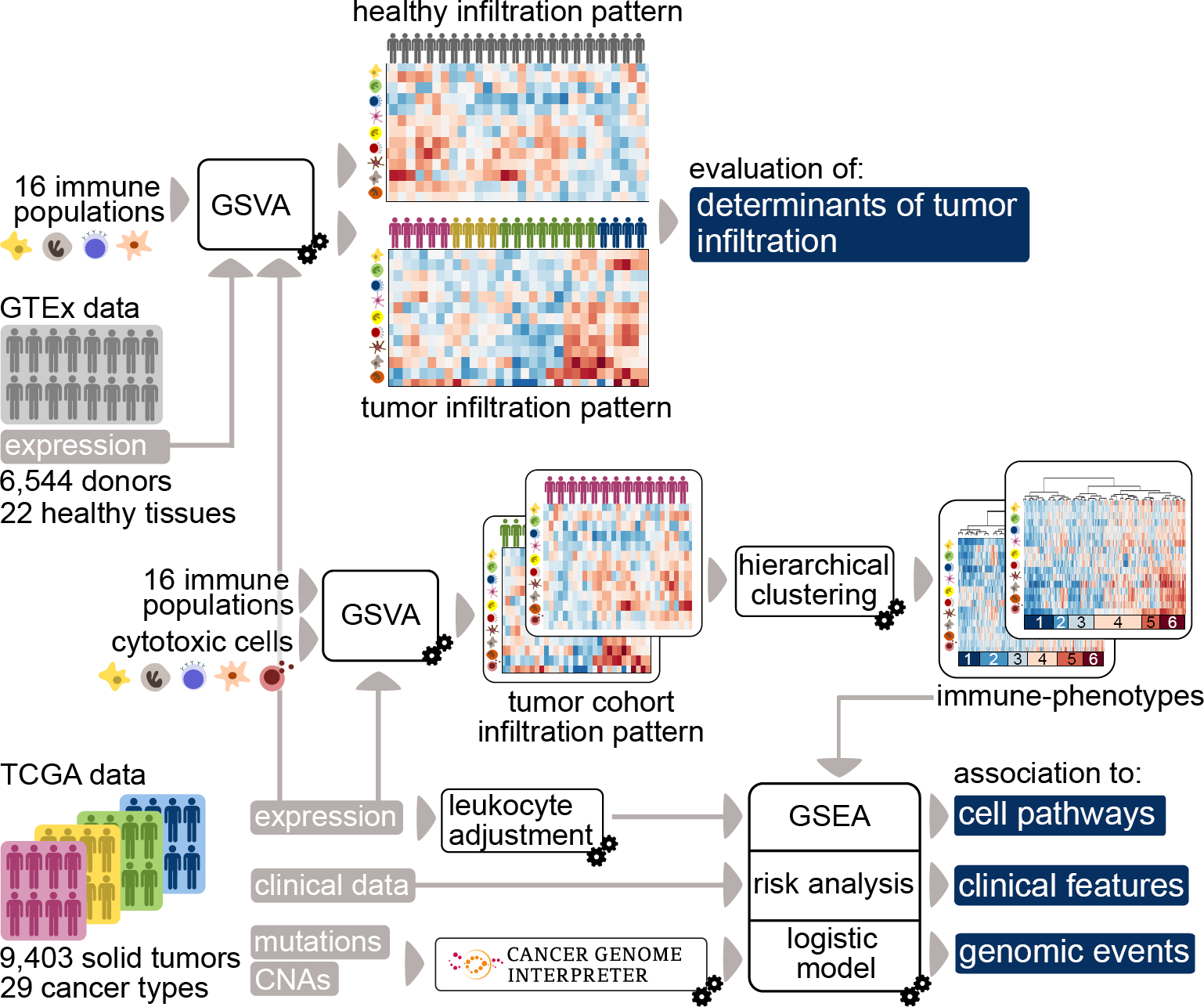
Overview of all analyses carried out in the study. We defined the immune infiltration pattern of a pan-cancer TCGA cohort with the abundance of a comprehensive set of immune cell populations estimated via a gene set variation analysis (GSVA). We first aimed to evaluate the contribution of the tissue of origin and cancer type in the heterogeneity of the immune infiltrate of tumors and thus we also computed the immune infiltration pattern of a cohort of GTEx healthy donors (upper part of the figure). Next, we sought to identify traits of the tumors that may influence (or be influenced) by the characteristics of the immune infiltrate. To this end, we computed cytotoxic-driven immune-phenotypes that clustered tumors on the basis of distinct selective pressures exerted by the immune infiltrate and identified clinical, genomic and transcriptomic traits associated to these scenarios (lower part of the figure) via an ensemble of computational analyses (GSEA: gene set enrichment analysis).

We first wanted to answer the question of how much of the infiltration pattern of tumors is determined by their tissue of origin or their cancer type, under the hypothesis that the remaining part could only be explained by the specific characteristics developed by each tumor. To this end, we estimated the immune infiltration pattern of 9,403 tumors from 29 solid cancers (Table 1) and compared it to the immune infiltration pattern of samples of 22 matching healthy tissues from 6,544 donors. We also computed the heterogeneity of the immune infiltration pattern of tumors across and within cancer types.

**Table 1.**

| Cancer type acronym | Cancer type full name | Number of patients (RNA-seq) |
| --- | --- | --- |
| ACC | Adrenocortical carcinoma | 78 |
| BLCA | Bladder Urothelial Carcinoma | 404 |
| BRCA | Breast invasive carcinoma | 924 |
| BRCA-TN | Breast triple negative invasive carcinoma | 158 |
| CESC | Cervical squamous cell carcinoma and endocervical adenocarcinoma | 301 |
| CHOL | Cholangiocarcinoma | 36 |
| COREAD | Colorectal adenocarcinoma | 599 |
| ESCA | Esophageal carcinoma | 182 |
| GBM | Glioblastoma multiforme | 163 |
| HNSC | Head and Neck squamous cell carcinoma | 515 |
| KICH | Kidney Chromophobe | 65 |
| KIRC | Kidney renal clear cell carcinoma | 515 |
| KIRP | Kidney renal papillary cell carcinoma | 285 |
| LGG | Brain Lower Grade Glioma | 514 |
| LIHC | Liver hepatocellular carcinoma | 368 |
| LUAD | Lung adenocarcinoma | 511 |
| LUSC | Lung squamous cell carcinoma | 485 |
| MESO | Mesotelioma | 87 |
| OV | Ovarian serous cystadenocarcinoma | 300 |
| PAAD | Pancreatic adenocarcinoma | 156 |
| PCPG | Pheochromocytoma and Paraganglioma | 178 |
| PRAD | Prostate adenocarcinoma | 493 |
| SARC | Sarcoma | 254 |
| SKCM | Skin Cutaneous Melanoma | 434 |
| STAD | Stomach adenocarcinoma | 405 |
| THCA | Thyroid carcinoma | 499 |
| UCEC | Uterine Corpus Endometrial Carcinoma | 357 |
| UCS | Uterine Carcinosarcoma | 57 |
| UVM | Uveal Melanoma | 80 |
| Total |  | 9403 |

After verifying that the heterogeneity of the infiltrate patterns observed in solid cancers cannot be explained solely by their tissue of origin or their cancer type, we set out to identify intrinsic tumor features that influence that pattern. We first grouped the tumors of each cancer type into clusters (immune-phenotypes), of similar immune infiltration pattern and level of cytotoxic cells. These immune-phenotypes represent qualitatively and quantitatively differentiated scenarios of immune infiltration and therefore, we reasoned that they could result in the selection of different mechanisms of evading the action of the immune system. To test this hypothesis, we systematically correlated the immune-phenotypes with: (i) clinical and pathological features of the tumors; (ii) genomic characteristics, including putative driver mutations and copy number alterations (identified by the Cancer Genome Interpreter^12^); and (iii) the activation status of cancer cell pathways (after subtracting the contribution of the immune cells to the expression of genes in the bulk sample). The following sections summarize the main results obtained in each of these steps.

### The immune infiltration pattern of tumors is heterogeneous across and within cancer types

We first sought to ascertain the extent to which the pattern of immune infiltration of tumors is driven by the tissue of origin and the cancer type. To answer this question, we computed the immune infiltration pattern –as the GSVA enrichment scores for 16 immune cell populations– across the pan-cancer cohort of 9,403 tumors (Table S2A). This analysis revealed that the relative abundance of many immune cell populations is correlated, indicating that they tend to co-infiltrate the tumor (Fig. S1). We then computed the healthy immune infiltration pattern of samples of 22 matching healthy tissues from 6,544 donors using the same method (Table S2B and Table S3). The immune infiltration pattern of tumors clearly deviated from that of their matched tissue of origin across cancer types (Fig. S2). At one end of the spectrum, LUSC, UCS, LUAD and BLCA presented the greatest number of immune cell populations with lower abundance than their tissues of origin. At the other end, PAAD, KIRC, GBM and SKCM possess the greatest number of immune populations with higher infiltration than their tissues of origin.

Next, we wanted to determine the influence of the type of cancer on the immune infiltration pattern. While the immune infiltration varies between malignancies (Figure S3), we reasoned that specific features of the tumor samples could contribute some degree of heterogeneity to the infiltration pattern within each cancer cohort. To test this hypothesis, we grouped the 9,403 tumors via a hierarchical clustering of their immune infiltration pattern (the GSVA enrichment scores of the 16 cell populations). We obtained 17 groups (Fig. 2A, see Methods), which reflect distinct ensembles of immune cells infiltrating solid tumors (Fig. S4). We then evaluated the heterogeneity of the infiltrate among tumors of each cancer type. We observed that tumors of most cancer types are distributed across multiple clusters (Fig. 2B). STAD, CHOL, LUSC, BRCA, BRCA-TN, SKCM and COREAD are the tumor types with the most heterogeneous infiltration landscape, whereas PRAD, PCPG, LGG and GBM exhibit the most homogeneous infiltrate, with the majority (>65%) of their samples grouped in a single cluster. We then explored how similar is the infiltration pattern of individual tumors of different cancer types. We observed that most clusters grouped tumors of different cancer types. Cluster 10, which is formed almost entirely by PRAD samples, is the most outstanding exception. Even tumors arising from tissues with very particular infiltration characteristics, such as the immune-privileged brain and eye^13,14^, clustered together with tumors that originate in other tissues. For example cluster 13 contains a high proportion of LGG, PCPG, KICH and CHOL tumors; cluster 7, an important fraction of GBM and THCA tumors; and cluster 14 has an important proportion of UVM, UCS, UCEC, BLCA, COREAD and SKCM tumors.

**Figure 2.**
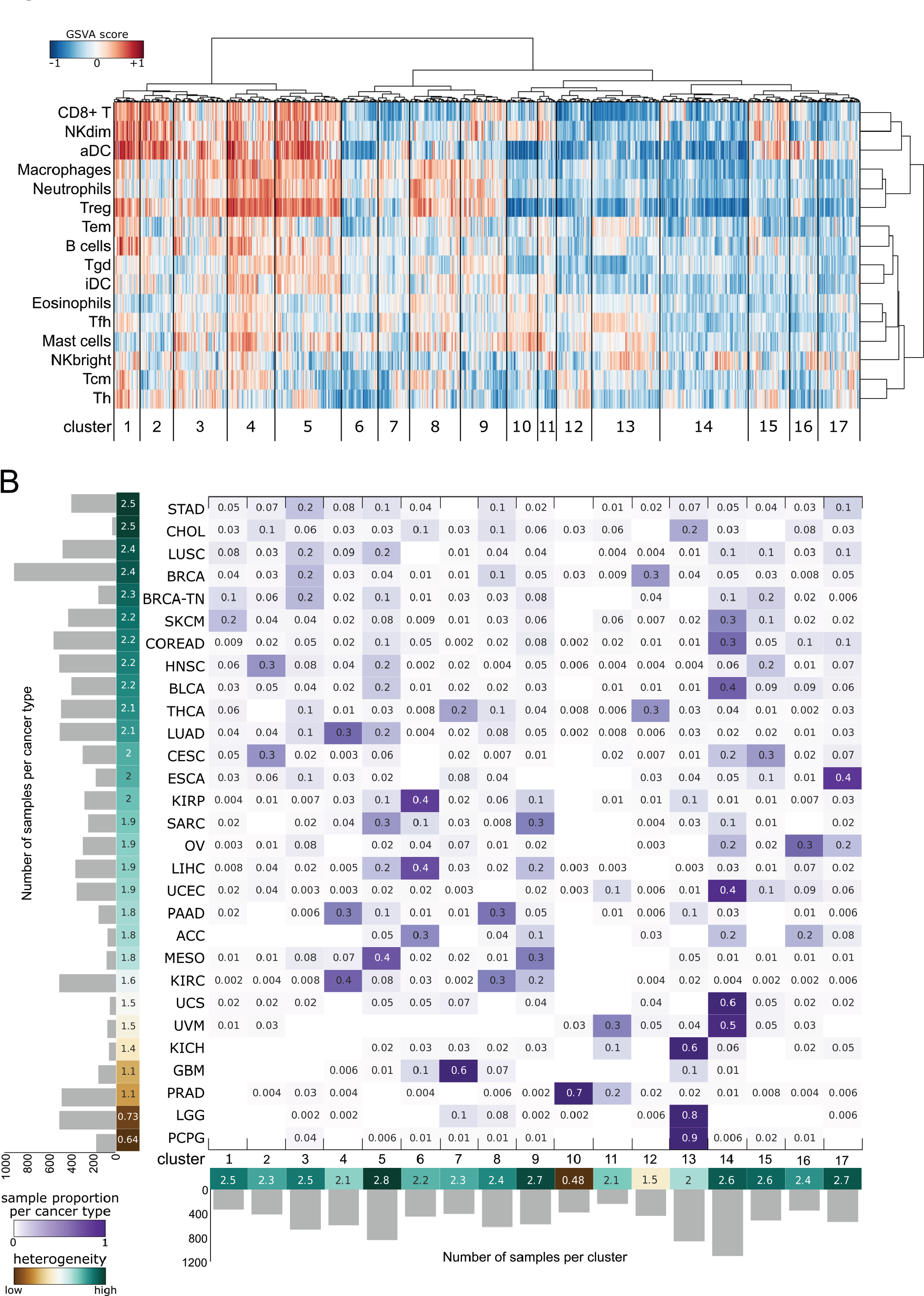
Determinants of the immune infiltration pattern. (A) Immune infiltrate clusters of 9,403 tumors of 29 solid cancers obtained through hierarchical clustering of the GSVA enrichment scores computed for 16 immune cell populations. We defined 17 groups (see Methods) that represent distinct patterns of immune infiltration of solid tumors. (B) Distribution of tumors of each malignancy across clusters of immune infiltration pattern. Numbers in the matrix represent the proportion of tumors of each cancer cohort in each cluster. A metric based on the entropy score (see Methods) represents how the tumors of each cancer type distribute across clusters (left bar) and how the tumors of different cancer types contribute to each cluster (bottom bar).

In summary, we observed that the pattern of immune infiltration of solid tumors differs from that observed in their tissue of origin. Furthermore, although the cancer type is an important determinant of the global infiltration of tumors, the infiltration pattern across cancer types is heterogeneous, and tumors of different malignancies show similar infiltration patterns. Thus, we conclude that specific features developed by each tumor could play an important role in shaping the immune infiltration pattern.

### Identification of various immune-phenotypes across cancer types

We then set out to identify which specific features of solid tumors –in combination with their tissue of origin and cancer type– contribute to define their immune infiltration pattern. We reasoned that different features may appear in tumors of different cancer types in response to immune infiltration patterns with varying cytotoxic capabilities. Therefore, we recomputed the immune infiltration pattern of tumors within each cancer cohort and used, as a proxy of the cytotoxic capability of the infiltrate, an additional gene signature representing all cytotoxic cells in the infiltration pattern (see Methods). As expected, GSVA scores of cytotoxic cells across tumors strongly correlated with those of CD8+ T and NKdim cells, whereas their correlation with Tgd cells was moderate. Interestingly, correlation between cytotoxic cells and NKdim cells was weaker across PRAD, LGG, GBM and ACC cohorts, which may suggest a smaller contribution of the innate immune system in the cytotoxicity of the infiltrate of these tumors (Fig. S5).

We grouped the tumors of each cancer cohort using a hierarchical clustering (Tables S2C to S2ZE) in which the GSVA score of the cytotoxic cells was given more weight than that of the other immune cell populations. Specifically, 25% of the total contribution to the clustering (see Methods) was given to this population. As a result, we grouped the tumors of each cohort into six (see Methods) cytotoxic-driven clusters (Fig. 3A, Fig. S6, Table S4). These clusters reflect different cases of immune infiltration pattern within each cancer type, regardless of differences between malignancies (Fig 3B). We call these clusters of tumors with similar infiltration pattern cancer immune-phenotypes. Although driven by the cytotoxic activity, the immune-phenotypes also reflect the the abundance of all individual immune cell populations. Immune-phenotypes range from an extreme scenario of poor overall immune infiltration and very low cytotoxic component (immune-phenotype 1 of each cancer type) to high relative abundance of most individual immune cell populations and cytotoxicity (immune-phenotype 6). Interestingly, the ratio between the abundance of *bona fide* immune effector (e.g. CD8+ T and Nkdim) and immune suppressive (e.g. macrophages, neutrophils, iDCs and Tregs) populations across immune-phenotypes grows significantly with the immune-phenotypes in most cases (Fig. S7). In summary, an increase of both the quantity and the quality of the immune infiltrate is observed from immune-phenotype 1 to 6.

**Figure 3.**
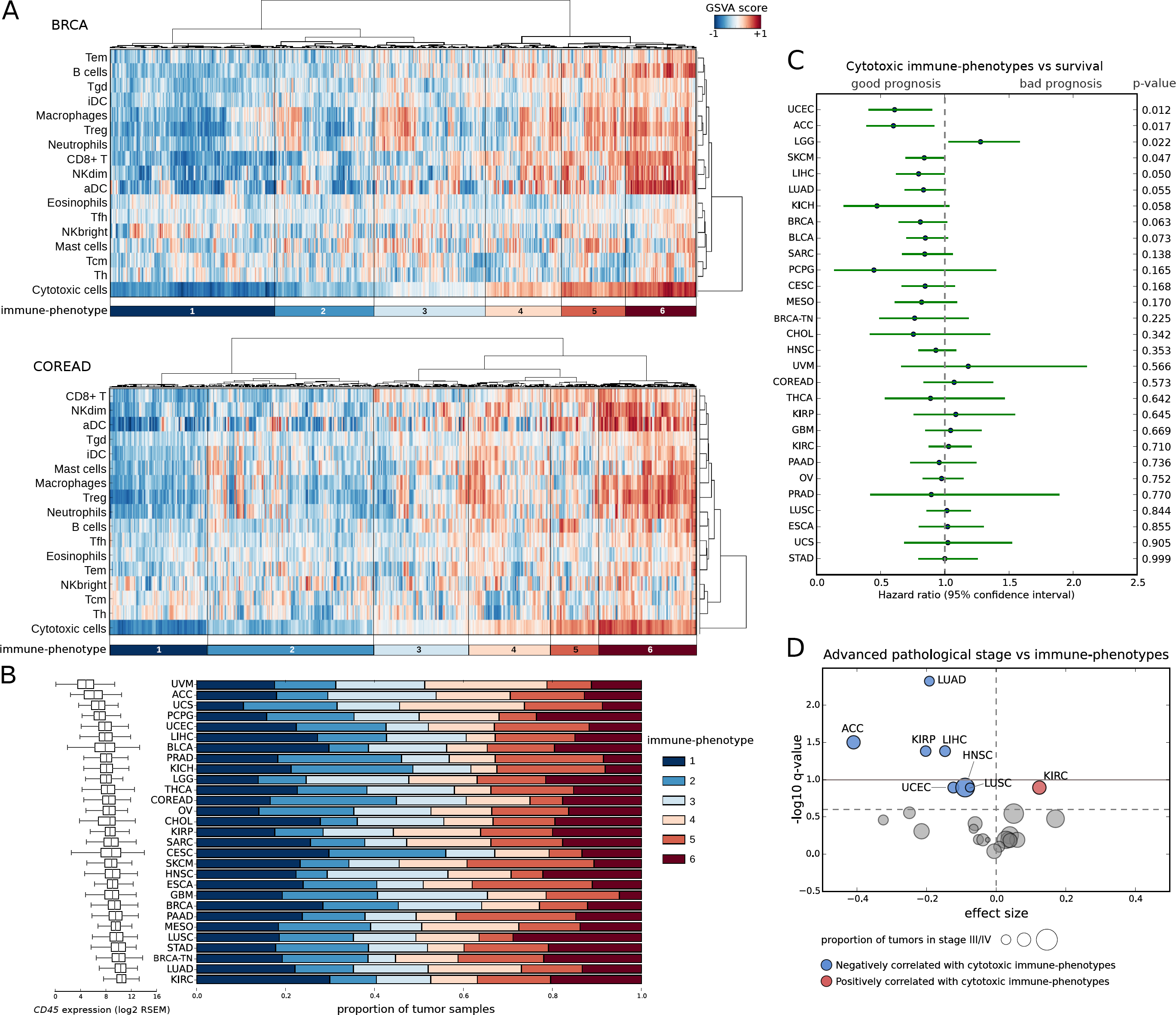
Tumor immune-phenotypes. (A) Cytotoxic-driven hierarchical clustering of the immune infiltration pattern of tumors yields six immune-phenotypes (see Methods) in each cancer cohort. The configuration of immune-phenotypes of the breast (BRCA; n=924) and colorectal (COREAD; n=599) cancer cohorts (the two largest) are displayed as illustration. Immune-phenotypes are numbered in ascending order of the median GSVA enrichment score of cytotoxic cells, and colored from dark blue to dark red. (B) Proportion of tumors with each immune-phenotype across cancer types. Note that immune-phenotypes represent tumors with distinct immune infiltrate settings within each cancer cohort regardless of the overall differences between malignancies. Cancer types are displayed in ascending order of their absolute leukocyte content (measured by the *CD45* expression of the bulk samples; left boxplot). (C) Influence of the immune-phenotypes on the overall survival. A hazard ratio lower than one indicates that patients with tumors of higher cytotoxic immune-phenotypes show improved survival. P-values were calculated with the Cox regression model. (D) Association between advanced (III/IV) pathological stage of the tumors and their immune-phenotypes at diagnosis. Significant events (linear regression q-value<0.2; see Methods) are depicted in red (associated with higher cytotoxic immune-phenotypes) and blue (associated with lower cytotoxic immune-phenotypes). Horizontal lines indicate q-values of 0.1 (solid line) and 0.2 (dot line).

We next explored whether these immune-phenotypes associated with clinical and pathological characteristics of the tumors. In coherence with previous reports^8,15^, we observed that the survival of patients improved with the increase of the cytotoxic level of the immune-phenotypes in several cancer types (Fig. 3C). An intriguing exception is LGG, the clinical outcome of which is worse for tumors with immune-phenotypes of higher cytotoxicity. This finding can be explained by the loss of integrity of the blood-brain barrier, which facilitates immune infiltration but also correlates with tumor size and aggressiveness^16,17^ (Fig. S8). Finally, we also observed more advanced pathological stage of tumors at diagnosis associate with higher cytotoxicity of the immune-phenotypes in nine cancer types. Renal clear cell carcinomas (KIRC), which show the opposite association, are an exception (Fig. 3D). This observation suggests that tumors preferentially progress in the presence of a poor cytotoxic immune infiltrate. Without longitudinal samples, however, it is not possible to distinguish whether this poorly cytotoxic immune-phenotype exists from the onset of the disease, or arises as a consequence of its development. In contrast, tumors with highly cytotoxic immune-phenotype would be partially kept in check by the immune system and progress less frequently to more advanced stages.

In summary, we identified cancer immune-phenotypes, which delineate different scenarios of quantity and quality of the immune infiltrate of solid tumors associated with clinical and pathological features.

### Different tumor genomic events appear across immune-phenotypes

We reasoned that different selective pressures reflected by diverse immune-phenotypes could elicit distinct tumoral mechanisms of immune surveillance escape. Therefore, we aimed to identify the genomic features of tumors that are associated with these immune-phenotypes.

First, we evaluated the coding mutation burden, known to be particularly high in certain tumors due to the underlying mutational processes^18^ and which has been shown to correlate with a strong anti-tumor immune response^19^. We focused on missense mutations and short frameshift indels, which are more likely to give rise to *de novo* antigens. We found a significant positive correlation between the coding mutational burden and the level of cytotoxicity of the immune-phenotype in five cancer types; however, a negative correlation was apparent in three other cancer types (Fig. 4A). Hypermutated tumors (defined on the basis of their relative mutational burden; see Methods) are significantly enriched for highly cytotoxic immune-phenotypes (immune-phenotypes 5 and 6) in COREAD, BRCA and STAD (odds ratio of 3.7, 3.4 and 2.1; Fisher’s *p* value <0.05 in all cases). Consistently, mutations most likely causing defective DNA repair mechanisms (signatures reflective of altered *POLE*, *BRCA-1/2* and mismatch repair deficiency) are significantly overrepresented in tumors with immune-phenotypes of higher cytotoxicity (Fig. S9). In agreement with a recent study^20^, we also observed that the burden of gene copy number alterations (CNAs) of tumors negatively correlated with the cytotoxicity of the immune-phenotypes in eleven cancer types (Fig. 4B). The clonal heterogeneity of tumors (estimated via the distribution of the variant allele frequency of their mutations^9^; see Methods) also negatively correlated with the cytotoxicity of the immune-phenotypes across twelve cancer types (Fig 4C). This may be the result of tumor immune-edition, in which the action of a highly cytotoxic immune infiltrate restricts the clonal diversity of tumors (even those with higher mutation burden).

**Figure 4.**
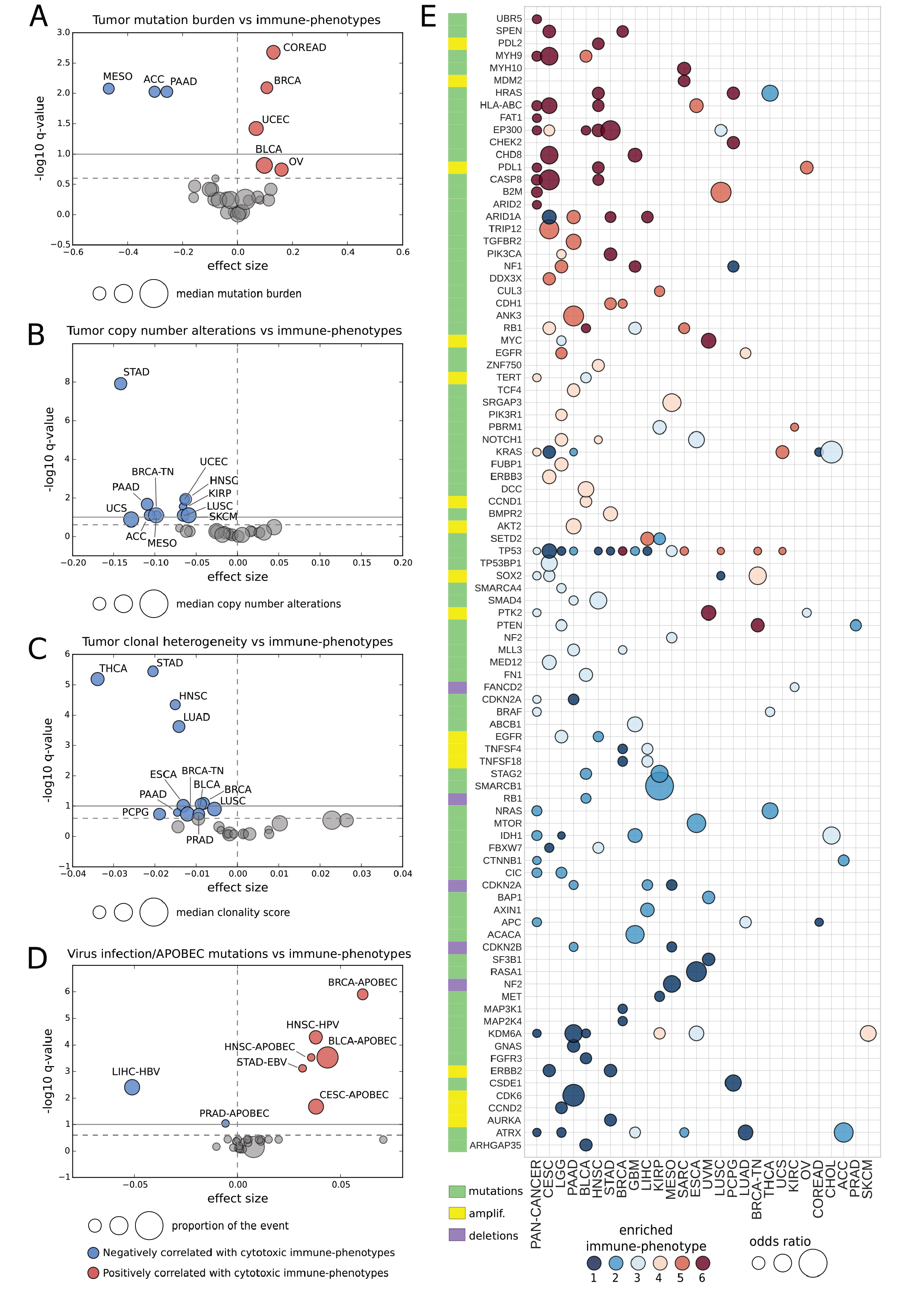
Tumor genomic features associated to immune-phenotypes. Association between the mutational (missense and frameshift) burden (A), the copy number alterations burden (B), the clonal heterogeneity (C) and the virus infection (D) of tumors and their immune-phenotypes. For the latter, the presence of known oncogenic viruses (HPV: human papillomavirus; EBV: Epstein-Barr virus; HCV: hepatitis C virus; HBV: hepatitis B virus; see Fig. S14 for the analysis of HPV types) and APOBEC mutational signatures were evaluated. Significant events (linear regression q-value<0.2; see Methods) are depicted in red (associated with highly cytotoxic immune-phenotypes) and blue (associated with poory cytotoxic immune-phenotypes). Horizontal lines indicate q-values of 0.1 (solid line) and 0.2 (dot line). (E) Genes with driver alterations significantly associated to each immune-phenotype (see Methods). Both cancer cohort-wise and pan-cancer analyses (see Methods) were carried out and significance was considered for q-values <0.1. Per-cancer associations with uncorrected p-value <0.05 that appeared significant in the pan-cancer analysis are also shown (HLA-A,-B and -C mutations are considered together). The color of the circle represents the cancer immune-phenotype significantly enriched by the gene driver alterations (see legend at the bottom), and the size is proportional to the magnitude of that enrichment. The type of driver alterations (mutations, amplifications or deletions) is color-coded at the left of the gene name. Cancer types are sorted in decreasing order of number of genes showing significant associations (at least one).

Viral infections are known to elicit an immune response driven by the presentation of viral antigens^19^. Therefore, we next asked whether the presence of viral infection in tumors (see Methods) is associated with specific immune-phenotypes. As a result, we found that the proportion of likely APOBEC generated mutations – often a consequence of viral infection^21^– positively correlates with the level of cytotoxicity of the immune-phenotypes in four cancer types (Fig. 4D and Fig. S9). Furthermore, the number of Epstein-Barr virus infected STAD and human papillomavirus infected HNSC tumors also positively correlate with the cytotoxicity of the immune-phenotypes, whereas no association between human papillomavirus infection and the immune-phenotype was apparent in CESC. Hepatitis C-infected hepatocellular carcinomas are enriched for highly cytotoxic immune-phenotypes, although the association did not reach statistical significance. Strikingly, hepatitis B-infected hepatocellular carcinomas are significantly overrepresented amongst those with poorly cytotoxic immune-phenotypes (Fig. 4D).

Finally, we focused on individual cancer driver genomic alterations (identified in each tumor of the 29 cohorts using the Cancer Genome Interpreter^12^) that significantly associate with the various immune-phenotypes. To address the statistical issues derived from the sparsity of the data, we implemented a regularized logistic regression analysis adjusting for the number of mutations and CNAs (see Methods). We regressed the genomic alterations on the immune-phenotypes of each cohort separately. In addition, we carried out the analysis in a pan-cancer manner by considering together the tumors with equivalent immune-phenotypes (all tumors with immune-phenotype 1 across all cohorts, all with immune-phenotype 2, and so on) to increase the statistical power to detect associations that are coherent across cancer types (Fig. 4E, Table S5). Some driver alterations significantly overrepresented among tumors with highly cytotoxic immune-phenotypes (immune-phenotypes 5 and 6) have already been described to elicit resistance to immune destruction. These include mutations of members of the *HLA* gene family and *B2M* (part of the antigen-presentation machinery), mutations of *CASP8* (extrinsic apoptosis pathway) and amplifications of the *PDL-1/2* genes (negative immune checkpoints)^3,4,7,22^. We also identified other genomic alterations significantly overrepresented among these tumors affecting epigenetic regulators (e.g. *EP300*, *ARID1A* and *ARID2*), E3 ubiquitin ligases (e.g. *UBR5*, *CUL3* and *TRIP12*) and genes encoding cell-cell interaction proteins (e.g. *MYH9*, *MYH10*, *ANK3*, *CDH1* and *FAT1*), all of which constitute novel potential drivers of immune resistance. At the opposite end of the spectrum, we identified several genes with alterations significantly overrepresented among tumors with poorly cytotoxic immune-phenotypes (immune-phenotypes 1 and 2). These include genes involved in the regulation of cell cycle, DNA replication and telomere maintenance (e.g. *ATRX, STAG2, AURKA, BAP1, FBXW7, CDKN2A, CDKN2B, CCND2* and *CDK6*), as well as genes of the WNT–β-catenin signaling pathway (e.g. *CTNNB1, APC, AXIN1*, and *SMAD4*). Of note, driver alterations affecting one gene are in some cases associated with distinct immune-phenotypes depending on the cancer type. Mutations of *NRAS/HRAS*, for example, associate with highly cytotoxic immune-phenotypes in HNSC and PCPG but to poorly cytotoxic immune-phenotypes in THCA. *NF1* mutations associate with highly cytotoxic immune-phenotypes in LGG and GBM, but with poorly cytotoxic immune-phenotypes in PCPG. *TP53* alterations, which significantly associate with various immune-phenotypes across thirteen malignancies constitute the most salient example of this behavior. Of note, no association between these driver alterations and individual immune cell populations explain these seemingly contradictory observations across cancers (Table S6).

Summing up, we identified genomic events that strongly associate with specific tumor immune-phenotypes, which constitute a catalog of putative genomic drivers of immune evasion. Some of these associations vary depending on the cancer type, which may be explained by the involvement of these genes in different cell processes in each malignancy. Therefore, we reasoned that a more complete picture of the mechanisms that tumors develop while progressing in these microenvironments can be achieved by analyzing the differential activity of their biochemical pathways.

### Several transcriptional programs appear over-activated across immune-phenotypes

We reasoned that cellular programs over-activated in tumors may differ depending on their immune-phenotype. To test this hypothesis, we selected a comprehensive collection of cancer cell pathways with minimal overlap between their gene sets (see Methods, Fig. S10 and Table S7). Then, we inferred their differential activation across immune-phenotypes from the collective up-regulation of the genes integrating them via a gene set enrichment analysis (GSEA)^23^. Because tumor samples are admixtures of tumor cells and their microenvironment, we applied the GSEA after adjusting the expression level of each transcript for the contribution of the immune content in the bulk sample (see Methods). This step prevented the overestimation of the contribution of tumors to the expression of genes that are highly expressed in immune cells (e.g. inflammatory cytokines or immune checkpoints) or the underestimation of genes whose expression is specific of tumors (e.g. cancer testis antigens) (Fig 5A, Fig. S11). Cancers with no matching healthy tissue data available in GTEx (CHOL, MESO and UVM) were excluded from the GSEA analysis.

**Figure 5.**
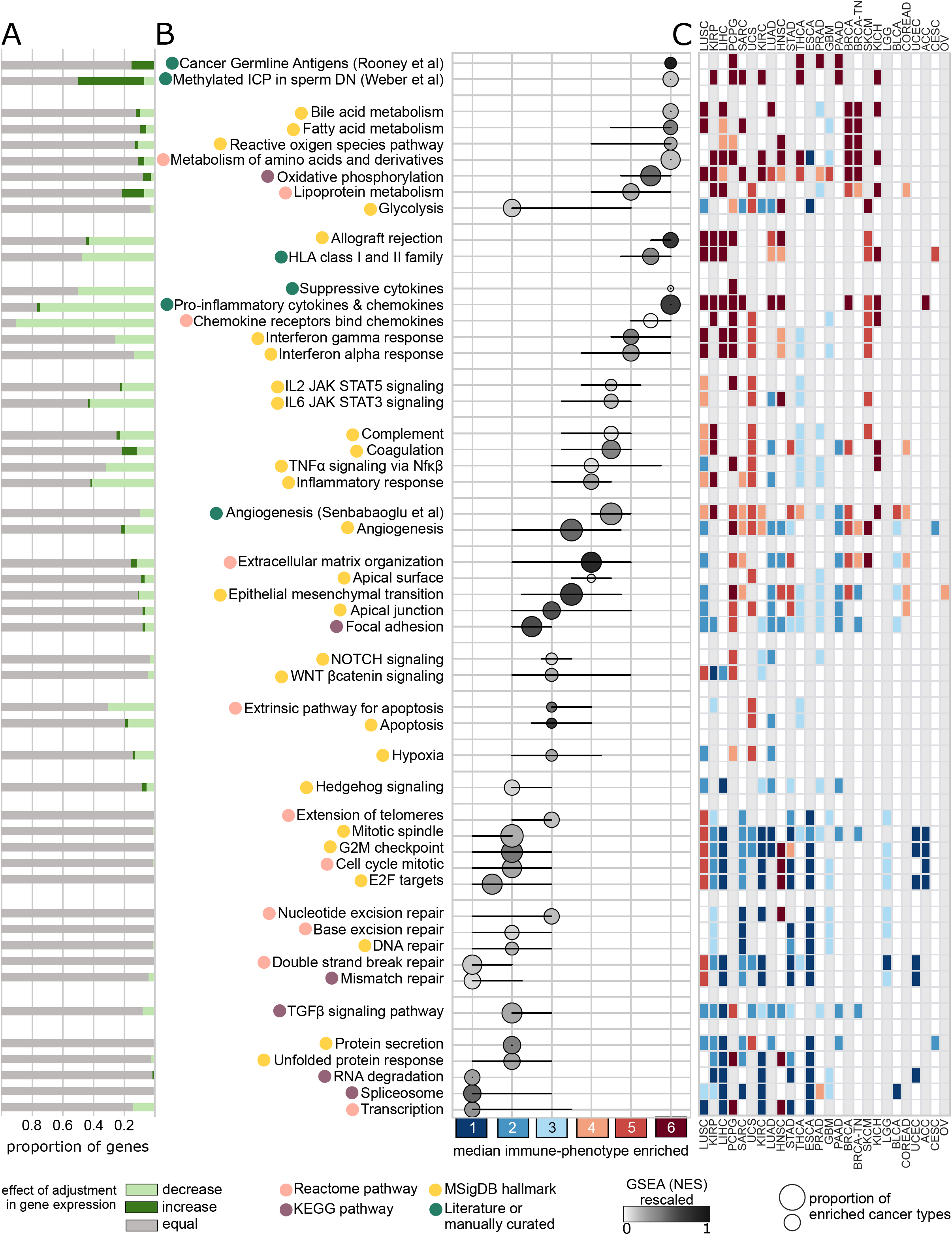
Transcriptional programs over-activated across immune-phenotypes. (A) Barplot of the proportion of genes in each pathway whose expression change after adjustment for the immune content of the tumor sample (pale green: decrease, dark green: increase, gray: unchanged; see Methods). (B) Over-activated pathways significantly enriched (q-value of the GSEA<0.25; see Methods) for tumors of each immune-phenotype. Pathways are grouped in the panel according to their broad cell processes. The position of each circle represents the median immune-phenotype (between 1 and 6) with significant enrichment for over-activation of the pathway across the 29 cancer types (black whiskers display percentiles 25 and 75). The size of the circles represents the number of cancer types in which these significant associations are observed, with the shade of the gray proportional to the median of the GSEA enrichment scores. (C) Heatmap detailing the immune-phenotype significantly enriched for the over-activation of each pathway across cancer types (see panel B). Cancers are sorted in the matrix according to the overall number of pathways that exhibit these significant associations.

Tumors with different immune-phenotypes present distinct sets of over-activated cellular programs (Fig. 5B and C, Table S8). The over-activation of several pathways that may be responsible of an increased presence of anti-tumor cells in the infiltrate is overrepresented among tumors with highly cytotoxic immune-phenotypes (immune-phenotypes 5 and 6). These potentially immunogenic signals include the cancer germline antigens, pro-inflammatory cytokines and chemokines and genes of the HLA I/II families. Tumors with highly cytotoxic immune-phenotypes are also enriched for the up-regulation of several pathways of the energy metabolism, with the exception of glycolysis. This observation is in line with recent studies that have shown that a metabolic competition is established between tumors and immune cells. The differentiation of the latter into cytotoxic populations relies on the availability of glucose –over other energy sources– in the microenvironment, which is hindered by tumors with over-active glycolysis^24,25^. Other pathways that appear up-regulated in this scenario, such as the JAK/STAT signaling or a set of suppressive cytokines may directly counteract the attack of a highly cytotoxic infiltrate^2–4^. As expected, the over-expression of the *PDL-1/PDL-2* negative immune checkpoints is also enriched among tumors with highly cytotoxic immune-phenotypes in most of the cancer types^4,19^ (Fig. S12).

The over-expression of pathways and processes related to tumor invasion and remodeling of neighboring tissues –response to hypoxia, angiogenesis, glycolysis, focal adhesion, inflammation, epithelial to mesenchymal transition and extracellular matrix remodeling– is overrepresented amongst tumors with immune-phenotypes of mid cytotoxicity (immune-phenotypes 3 and 4). Their over-expression, however appears spread across several immune-phenotypes, which could be an indication that these tumors represent a transitional state between those with highly and poorly cytotoxic immune-phenotypes. Finally, poorly cytotoxic immune-phenotypes (immune-phenotypes 1 and 2) are enriched for up-regulated TGFβ and Hedgehog signaling pathways, which have been described to prevent immune infiltration via several mechanisms^26,27^. Besides, these immune-phenotypes are also strongly associated with the over-expression of genes involved in the progression of the cell cycle, RNA and protein synthesis, response to DNA damage, and telomere maintenance. The over-expression of this set of processes may be the result of tumors subject to an accelerated rate of growth and proliferation, which preferentially occurs in the presence of a low effective immune infiltrate^28,29^.

## DISCUSSION

Here we describe a comprehensive pan-cancer landscape of immune infiltration patterns across 29 solid cancers and identify genomic and transcriptomic tumor features that are significantly associated with distinct immune-phenotypes. As the computational estimation of the immune infiltration pattern of tumors presents several limitations, we have combined the use of robust mathematical models and bioinformatics methods aimed at refining the analysis, such as the identification of driver genomic events and the adjustment of expression of genes to account for the immune content of each tumor sample. The systematic analysis of thousands of tumors increases the statistical power to detect genomic and transcriptomic features that strongly associate with the events of interest. Finally, and differently from other studies focused on particular cancer types, our analysis identifies associations between tumor features and immune cells that appear consistently across various malignancies.

The immune infiltration pattern is heterogeneous within and across tumor types. We observed that the characteristics of the immune infiltrate are shaped by both the type of cancer and traits of individual tumors. We therefore identified immune-phenotypes that represent distinct quantitative and qualitative scenarios of immune infiltration in solid tumors. On the basis of these scenarios and the tumor features that we found significantly associated with each of them, we propose a general model that –despite variations between cancer types (Fig. 6)– describes the progression of tumors in these diverse microenvironments.

**Figure 6.**
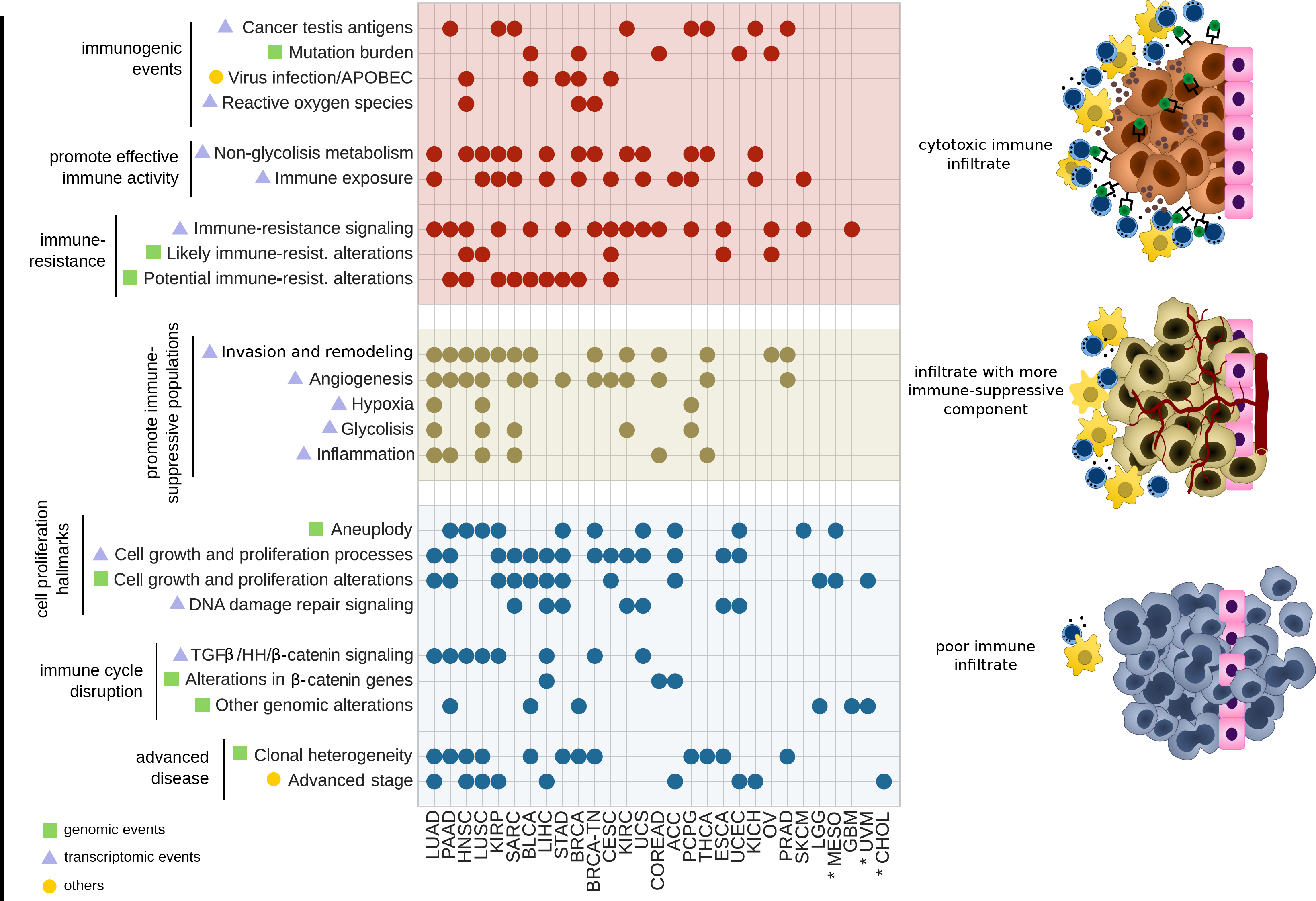
A model of tumor development under different scenarios of immune infiltration. The left panel summarizes tumor features associated to three scenarios of immune infiltration integrated by the tumors of immune-phenotypes 5-6 (highly cytotoxic, top), immune-phenotypes 3-4 (mid cytotoxic and a larger contribution of the immune-suppressive component, center), and immune-phenotypes 1-2 (poor overall immune infiltration, bottom). Each row groups several genomic or transcriptomic events (labeled as green squares and purple triangles, respectively) under broad terms (see Table S11). Circles mark the cancer types in which events included within the term are associated to one of the scenarios of immune infiltration (see Methods). Cancer types are sorted in descending order of number of overall significant associations. Cohorts marked with an asterisk (CHOL, UVM and MESO) do not contemplate associations with any transcriptomic program (due to the lack of matching healthy tissue expression to perform the leukocyte correction of the tumor samples data). TGFβ: transforming growth factor beta; HH: Hedgehog.

The first scenario describes the progression of tumors with highly cytotoxic immune-phenotype (Fig. 6, top), which receive the most robust attack from the host’s immune system. These tumors are enriched for immunogenic events, such as the over-expression of cancer testis antigens and virus infection.. They also carry a high mutational burden, frequently accompanied by fewer CNAs (Fig. S13) and associated with alterations of genes of the intrinsic DNA damage response pathway (Table S9). These tumors also show an increased activity of non-glycolytic metabolic pathways. While this may leave enough glucose in the microenvironment to maintain a cytotoxic immune population^24,25^, this can also result in high rates of oxidative damage to the DNA of these tumors. Therefore, the high cytotoxicity of their infiltrate, which may trigger the over-activation of the antigen-presentation machinery and the interferon signaling (immune exposure), may thus be partially explained by the increased number of neoepitopes and DNA damage^30^. In consequence, these tumors develop mechanisms to resist the immune assault, such as the over-expression of the JAK/STAT signaling and several inhibitory chemokines and negative immune checkpoints (immune-resistance signaling). They also acquire loss-of-function mutations of *HLA*, *B2M* and *CASP8*, and amplifications of *PDL-1/2*^*3,4,22*^. Other genomic alterations overrepresented amongst these tumors affect a variety of epigenetic factors, E3 ubiquitin ligases and cell-cell interaction genes involved in various tumor processes that may potentially drive resistance to the immune attack^31–36^.

In the second scenario (Fig. 6, center), tumors with immune-phenotypes of mid cytotoxicity are enriched for the over-expression of processes involved in the invasion and remodeling of neighboring tissues, such as the epithelial to mesenchymal transition, focal adhesion, and extracellular matrix remodeling. They are also enriched for the over-expression of other hallmarks of tumor progression, such as angiogenesis, glycolisis, inflammation and hypoxia. Of note, these processes are extensively associated with the infiltration of immune suppressive populations^24,25,37–40^. Coherently, we observed that the immune infiltration pattern of tumors of this group displays a shift towards the immune-suppressive component. Tumor cells that progress in this scenario may thus –at least partly– keep the immune surveillance in check through mechanisms that foster the recruitment of an immune-suppressive infiltrate.

The third scenario describes the progression of tumors with poorly cytotoxic immune-phenotypes (Figure 6, bottom), which present high aneuplody (partially explained by the occurrence of TP53 mutations in some cancer types; Table S10). They also over-express – and/or present genomic alterations of– genes involved in key processes for cell growth and proliferation, such as the regulation of cell cycle, cell division, telomere maintenance and mRNA and protein synthesis. Some of these molecular traits may contribute to the disruption of the cancer-immunity cycle, and thus drive immune evasion, while others may constitute opportunistic events favored by the poorly cytotoxic immune infiltrate. Examples of the former are the mutations of *IDH1* and WNT-β-Catenin genes and the over-activation of the TGFβ and Hedgehog signaling pathways, all of them reported to reduce immune cell infiltration^26,27,41,42^. These tumors also exhibit worse prognosis in several cancer types which is partially explained by the faster progression of the disease (measured by their pathological stage) in such a permissive infiltrate. The lower anti-tumor pressure exerted by these immune-phenotypes also allows the appearance of more tumor clones, resulting in more heterogeneous malignancies with greater metastatic potential^43^.

It is not clear from our results to what point these three scenarios represent tumors with separate evolutionary pathways or different stages in their progression. In the latter case, tumors with highly cytotoxic immune-phenotype would be at early stages, and equipped with mechanisms to resist a robust immune attack. They would then progressively evolve to invade neighboring tissues while their immune-phenotype shifts towards a more immune-suppressive infiltration pattern. Tumors with a poor cytotoxic infiltrate would represent advanced stages of a malignancy in full progression and virtually unchecked by the host’s immune system.

The most immediate result of our analysis is the identification of tumor features that are positively selected due to their interaction with the immune infiltrate. In this regard, we describe a general approach to classify tumors on the basis of their immune-phenotype. More long-term outcomes of this work involves understanding how these somatic genomic events and transcriptomics programs actively shape the immune infiltrate. This knowledge shall have important implications for the design of novel therapeutic approaches^31^. Currently, the most widely exploited immunotherapies involve immune checkpoint inhibitors. We explored transcriptomics data of small cohorts of patients with metastatic melanomas that received anti-CTLA4 and anti-PD1 therapies (42 and 28 patients, respectively)^44,45^. Patients with tumors with immune infiltration pattern with high cytotoxicity tend to respond better to the anti-CTL4 treatment. On the other hand, those with tumor with over-expression of gene sets involved in the WNT-β-catenin, extracellular matrix organization and angiogenesis pathways respond worse to anti-PD-1 (see Suppl. Note). Further analysis of larger cohorts of patients treated with immunotherapies will continue to clarify which tumor features are better predictors of their response. We also envision that the inclusion of drugs able to counteract mechanisms of immune-resistance or immune-suppression in clinical trials will reveal their potential for synergies with existing immunotherapies. Some of our findings, such as those supporting the use of drugs targeting the cell cycle to favor improvement of the quality of the immune infiltrate, or epigenetic modulators to improve the response to negative immune checkpoint inhibitors^28,32^, constitute promising examples to explore in this direction. In summary, tumor features strongly associated with distinct immune-phenotypes presented in this study may help interpret the response of solid tumors to immunotherapies and guide the development of novel drug combination strategies.

## METHODS

### Sample data collection and processing

TCGA data –mutations, copy number alterations, gene expression (RSEM gene-normalized), clinical annotations (tumor stage was manually annotated using the available pathologic and clinical data)– for 9,403 tumors across 29 cancer types was downloaded from the most recent freeze contained in the Firebrowse server (2016_01_28). The expression data (RPKM) of 8,304 donors across 22 healthy tissues was downloaded from the GTEx portal (version 6). The quantification of virus expression across tumors was retrieved from previous publications^7,46^. We considered a tumor infected if the expression of a virus was higher than the observed across healthy tissues, following the approach described by Rooney et al ^7^.

A tumor was considered hypermutated on the basis of its mutation burden as compared to the observed across that cancer type (the threshold was set to the median mutation burden of the cohort plus 4.5 times the interquartile range, minimum of 500 mutations). The clonal heterogeneity of each tumor was computed as the ratio between the dispersion of the variant allele fraction of the mutations in the sample and its median variant allele fraction ^9^. We analysed the RNA-seq data of samples from melanoma patients treated with anti-CTLA4^44^ (RPKM matrix provided by the authors, 42 patients) and anti-PD1^45^ (FPKM matrix downloaded from GEO: GSE78220, 28 patients) therapies.

### Estimation of the abundance of immune cell populations

Infiltration levels of sixteen immune populations were computed from the RNA-seq of each bulk sample. We selected immune cell populations that describe the infiltrate whose relative abundance can be computed with confidence from the expression data of the admixture samples given available gene sets form several sources (Suppl. Material). On detail, we used the gene set variation analysis (GSVA)^11^, which demonstrated to be more appropriate for our aim than available deconvolution-based methods (Suppl. Material). Immune populations were represented by non-overlapping gene sets specifically over-expressed in each cell type, selected from several previous publications^8,47,48^ (Suppl. Material and Table S1). We also calculated the infiltration levels of cytotoxic cells as a proxy of the anti-tumor activity of the overall immune infiltrate. The gene set to compute the abundance of cytotoxic cells includes genes over-expressed in CD8+ T, Tgd and NK cells (Suppl. Material). The GSVA produces normalized enrichment scores ranging from -1 to 1, which represent the abundance of the immune cell population in the sample relative to the remaining tumors in the analyzed cohort. Overall, thirty-one independent GSVA analyses were carried out for the pan-cancer cohort (all tumor samples pooled together, n=1), the healthy donors cohort (all healthy samples pooled together, n=1), and each cancer cohort separately (n=29).

### Comparison of the immune infiltration pattern of tumors and their tissue of origin

We first matched each solid cancer type in TCGA to its closest corresponding healthy tissue included in the GTEx project (Table S3). We then compared the GSVA scores computed for 9,403 tumors with those in matching healthy tissue samples (Suppl. Material). We considered that the abundance of a given immune population was different between tumors and healthy samples if the distribution of GSVA scores across tumors of the cancer type differed significantly from that measured across the matching healthy samples (Mann-Whitney q-value<0.05) and more that 0.2 of difference between their respective medians.

### Tumor samples clustering

Tumors were grouped in clusters of different immune infiltration pattern via a hierarchical agglomerative clustering (Euclidean distance and Ward’s linkage method). To cluster the tumors across the pan-cancer cohort (for the analysis described in the second section of Results), we used the GSVA scores of 16 immune cell populations of the 9,403 tumors pooled together. To cluster the tumors of each cohort separately (to build the immune-phenotypes), we included the GSVA scores of cytotoxic cells, and over-weighed this signature by a factor of four (Suppl. Material) in the process of clustering. The optimal number of clusters of tumors in each cohort was determined based on the percentage of variance of the data explained by them (Suppl. Material). In the analysis of the pan-cancer cohort, we used the changes in the distribution of cancer types across immune clusters as an additional criterion to choose this value (Suppl. Material).

### Quantification of mutational signatures

The deconstructSigs software^49^ was used to compute the contribution of previously reported mutational signatures (http://cancer.sanger.ac.uk/cosmic/signatures) to the overall landscape of mutations of each tumor. This calculation was limited to tumor samples with at least 50 mutations.

### Identification of genomic drivers associated to immune-phenotypes

We first identified genomic alterations (point mutations, small indels and gene copy number gains or losses) that are potential drivers of each tumor using the Cancer Genome Interpreter^12^ (https://www.cancergenomeinterpreter.org). Then, to identify the genes with driver alterations that significantly associate with a particular immune-phenotype, we implemented a regularized logistic regression analysis adjusted by the count of protein-affecting mutations and gene copy number alterations. Our model specifically addresses the spurious fitting derived from the sparsity of the data and provides empirical significance scores (Suppl. Material). The choice of regularization algorithm can be interpreted as a Bayesian logistic regression with a Gaussian weakly-informative prior distribution for the parameters of the immune clusters and adjustment covariates. We applied the logistic regression to each cancer cohort separately and we also performed a pan-cancer analysis in which all the tumors with the same immune-phenotype (the number representing the cytotoxicity level) were pooled together regardless of their cancer type. Only genes bearing mutations in at least 2.5% of the sample of the cohort and/or CNAs in at least 5% were included.

### Expression adjustment

To adjust the gene expression measured in a tumor bulk sample by its immune content, we followed the rationale described in^50^. Briefly, we adjusted the expression value of each gene in each tumor sample according to the contribution of *CD45* to the expression of that gene observed in the matching healthy tissue (Suppl. Material and Fig. S11). Cancers with no matching healthy tissue data available (CHOL, MESO and UVM) were excluded from this analysis.

### Pathway enrichment analysis

We identified over-expressed pathways among tumors of a certain immune-phenotype running a Gene Set Enrichment Analysis (GSEA)^23^ on the adjusted expression data (available for 9,200 tumors across 26 cancer types, due to the exclusion of the cohorts of CHOL, MESO and UVM; see above). Significance was considered for values of corrected *p* (which takes into account the size of each gene set plus the multiple test correction, as provided by the method) lower than 0.25, as recommended by the authors (http://software.broadinstitute.org/gsea/doc/GSEAUserGuideFrame.html?Interpreting_GSEA). Gene sets were downloaded from the MSigDB database^23^. We included broad hallmarks and specific pathways of interest from the curated gene sets/canonical pathways collection. In addition, we manually gathered pathways not found in the aforementioned collections. Overall, we considered a total of 51 pathways (Table S7) with minimal gene set overlap between them (Fig. S10A) as well as with the gene sets used to compute the immune cells abundance (Fig. S10B).

### General model of association between tumor features and immune-phenotypes

We grouped the genomic and transcriptomic features of the tumors studied here into several broad groups that summarize our results. For example, the transcriptomic term ‘*invasion and remodeling*’ comprises the findings for the epithelial to mesenchymal transition (MsigDB Hallmarks), extracellular matrix organization (Reactome database) and focal adhesion (KEGG database) pathways. The details of the mapping of pathways and genes to broad terms employed in Figure 6 are included in Table S11. To build this figure, we considered that each of these broad terms was associated to the highly cytotoxic immune scenario if the median of the immune-phenotypes (numbered from 1 to 6 as explained above) associated to all the features integrating the term was greater than 4.5, to the poorly cytotoxic immune scenario if the median was lower than 2.5, and to the mid cytotoxic immune scenario otherwise.

### Statistical tests

Unless explicitly stated, the association between pairs of categorical variables was evaluated using the Fisher’s exact test. The distributions of two sets of any continuous variable were compared using the Mann-Whitney U test. The homogeneity of the distribution of samples across different groups was computed via an entropy score (described in Supplementary Methods). The influence of the immune-phenotype on the survival was evaluated via the Cox proportional hazard model. A linear regression was used to assess whether the value of a variable (e.g. the mutation burden) increases (or decreases) across immune-phenotypes. P-values were corrected using the Benjamini-Hochberg False Discovery Rate method as appropriate.

## ACKNOWLEDGEMENTS

D.T. is supported by project SAF2015-74072-JIN, which is funded by the Agencia Estatal de Investigacion (AEI) and Fondo Europeo de Desarrollo Regional (FEDER). C.R-P. is supported by an FPI fellowship (BES-2013-063354) from the Spanish Ministry of Economy and Competitiveness. N.L-B. acknowledges funding from the European Research Council (consolidator grant 682398). A.G-P. is supported by a Ramón y Cajal contract (RYC-2013-14554). The results shown here are in whole or part based upon data generated by the TCGA Research Network. Data generated by the Genotype-Tissue Expression (GTEx) Project --supported by the Common Fund of the Office of the Director of the National Institutes of Health, and by NCI, NHGRI, NHLBI, NIDA, NIMH, and NINDS--was also used. We thank Elena Gros (PhD) and Ignacio Melero (MD PhD) for the scientific review and helpful discussions of the study, Fiorella Ruiz for the annotation of the TCGA clinical data, and Dvir Aran (PhD) for the support in the expression adjustment methodology. We also thank Eliezer Van Allen (MD PhD) for providing the transcriptomic data of melanomas patients treated with anti CTLA-4 therapy.

### AUTHORS CONTRIBUTION

D.T designed the study and A.G-P and D.T coordinated the study and the analysis of the results. C.R-P carried out the immune profiling of RNA-seq samples and the transcriptomic analysis. D.T. computed the immune-clusters and their correlation with clinical and genomic characteristics. D.T and C.R-P. drafted the manuscript and prepared the figures. D.T, C.R-P and AG-P prepared the final version of the manuscript. F.M. developed the logistic regression model and supervised the use of statistical methods. S.R. performed the mutation signature analysis. R.D, N.L-B, A.M and J.M.P provided scientific feedback and contributed to the discussion of results. All the authors reviewed and approved the content and results of the study.

## REFERENCES

1. Hanahan, D. & Weinberg, R. a. Hallmarks of cancer: the next generation. Cell 144, 646–74 (2011).

2. Gajewski, T. F., Schreiber, H. & Fu, Y.-X. Innate and adaptive immune cells in the tumor microenvironment. Nat. Immunol. 14, 1014–22 (2013).

3. Khong, H. T. & Restifo, N. P. Natural selection of tumor variants in the generation of ‘tumor escape’ phenotypes. Nat. Immunol. 3, 999–1005 (2002).

4. Sharma, P., Hu-Lieskovan, S., Wargo, J. A. & Ribas, A. Primary, Adaptive, and Acquired Resistance to Cancer Immunotherapy. Cell 168, 707–723 (2017).

5. Newman, A. M. & Alizadeh, A. A. High-throughput genomic profiling of tumor-infiltrating leukocytes. Current Opinion in Immunology 41, 77–84 (2016).

6. Senbabaoglu, Y. et al. Tumor immune microenvironment characterization in clear cell renal cell carcinoma identifies prognostic and immunotherapeutically relevant messenger RNA signatures. Genome Biol. 17, 231 (2016).

7. Rooney, M. S., Shukla, S. A., Wu, C. J., Getz, G. & Hacohen, N. Molecular and genetic properties of tumors associated with local immune cytolytic activity. Cell 160, 48–61 (2015).

8. Charoentong, P. et al. Pan-cancer Immunogenomic Analyses Reveal Genotype-Immunophenotype Relationships and Predictors of Response to Checkpoint Blockade. Cell Rep. 18, 248–262 (2017).

9. Karn, T., Jiang, T., Hatzis, C. & al, et. Association between genomic metrics and immune infiltration in triple-negative breast cancer. JAMA Oncol. (2017).

10. Ali, H. R., Chlon, L., Pharoah, P. D. P., Markowetz, F. & Caldas, C. Patterns of Immune Infiltration in Breast Cancer and Their Clinical Implications: A Gene-Expression-Based Retrospective Study. PLoS Med. 13, (2016).

11. Hänzelmann, S., Castelo, R. & Guinney, J. GSVA: gene set variation analysis for microarray and RNA-seq data. BMC Bioinformatics 14, 7 (2013).

12. Tamborero, D. et al. Cancer Genome Interpreter Annotates The Biological And Clinical Relevance Of Tumor Alterations. bioRxiv (2017).

13. Carson, M. J., Doose, J. M., Melchior, B., Schmid, C. D. & Ploix, C. C. CNS immune privilege: Hiding in plain sight. Immunological Reviews 213, 48–65 (2006).

14. Oliva, M., Rullan, A. J. & Piulats, J. M. Uveal melanoma as a target for immune-therapy. Ann. Transl. Med. 4, 172 (2016).

15. Senovilla, L. et al. Trial watch: Prognostic and predictive value of the immune infiltrate in cancer. Oncoimmunology 1, 1323–1343 (2012).

16. Provenzale, J. M., Wang, G. R., Brenner, T., Petrella, J. R. & Sorensen, A. G. Comparison of permeability in high-grade and low-grade brain tumors using dynamic susceptibility contrast MR imaging. AJR Am. J. Roentgenol. 178, 711–716 (2002).

17. Seitz, R. J. & Wechsler, W. Immunohistochemical demonstration of serum proteins in human cerebral gliomas. Acta Neuropathol. 73, 145–152 (1987).

18. Alexandrov, L. B. et al. Signatures of mutational processes in human cancer. Nature 500, 415–21 (2013).

19. Chen, D. S. & Mellman, I. Elements of cancer immunity and the cancer-immune set point. Nature 541, 321–330 (2017).

20. Davoli, T., Uno, H., Wooten, E. C. & Elledge, S. J. Tumor aneuploidy correlates with markers of immune evasion and with reduced response to immunotherapy. Science (80-.). 355, eaaf8399 (2017).

21. Koito, A. & Ikeda, T. Intrinsic immunity against retrotransposons by APOBEC cytidine deaminases. Frontiers in Microbiology 4, p(2013).

22. Zaretsky, J. M. et al. Mutations Associated with Acquired Resistance to PD-1 Blockade in Melanoma. N. Engl. J. Med. 375, 819–29 (2016).

23. Subramanian, A. et al. Gene set enrichment analysis: A knowledge-based approach for interpreting genome-wide expression profiles. Proc. Natl. Acad. Sci. 102, 15545–15550 (2005).

24. Chang, C.-H. & Pearce, E. L. Emerging concepts of T cell metabolism as a target of immunotherapy. Nat Immunol 17, 364–368 (2016).

25. Ho, P. C. & Liu, P. S. Metabolic communication in tumors: a new layer of immunoregulation for immune evasion. J. Immunother. Cancer 4, 1 (2016).

26. Pickup, M., Novitskiy, S. & Moses, H. L. The roles of TGFβ in the tumour microenvironment. Nat. Rev. Cancer 13, 788–99 (2013).

27. Hanna, A. & Shevde, L. A. Hedgehog signaling: modulation of cancer properies and tumor mircroenvironment. Mol. Cancer 15, 1 (2016).

28. Goel, S. et al. CDK4/6 inhibition triggers anti-tumour immunity. Nature 548, 471–475 (2017).

29. Vinay, D. S. et al. Immune evasion in cancer: Mechanistic basis and therapeutic strategies. Seminars in Cancer Biology 35, S185–S198 (2015).

30. Mouw, K. W., Goldberg, M. S., Konstantinopoulos, P. A. & D’Andrea, A. D. DNA Damage and Repair Biomarkers of Immunotherapy Response. Cancer Discov. 617– 632 (2017). doi:10.1158/2159-8290.CD-17-0226

31. Gotwals, P. et al. Prospects for combining targeted and conventional cancer therapy with immunotherapy. Nat. Rev. Cancer 17, 286–301 (2017).

32. Mazzone, R., Zwergel, C., Mai, A. & Valente, S. Epi-drugs in combination with immunotherapy: a new avenue to improve anticancer efficacy. Clin. Epigenetics 9, 59 (2017).

33. Newbold, A., Falkenberg, K. J., Prince, H. M. & Johnstone, R. W. How do tumor cells respond to HDAC inhibition? FEBS J. 283, 4032–4046 (2016).

34. Fulda, S., Rajalingam, K. & Dikic, I. Ubiquitylation in immune disorders and cancer: From molecular mechanisms to therapeutic implications. EMBO Molecular Medicine 4, 545–556 (2012).

35. Banh, C., Fugère, C. & Brossay, L. Immunoregulatory functions of KLRG1 cadherin interactions are dependent on forward and reverse signaling. Blood 114, 5299–5306 (2009).

36. Li, Y. et al. Structure of Natural Killer Cell Receptor KLRG1 Bound to E-Cadherin Reveals Basis for MHC-Independent Missing Self Recognition. Immunity 31, 35–46 (2009).

37. Condeelis, J. & Pollard, J. W. Macrophages: Obligate partners for tumor cell migration, invasion, and metastasis. Cell 124, 263–266 (2006).

38. Facciabene, A. et al. Tumour hypoxia promotes tolerance and angiogenesis via CCL28 and Treg cells. Nature 475, 226–230 (2011).

39. Kumar, V. & Gabrilovich, D. I. Hypoxia-inducible factors in regulation of immune responses in tumour microenvironment. Immunology 143, 512–519 (2014).

40. de Mingo Pulido, A. & Ruffell, B. in Advances in Cancer Research 132, 139–163 (2016).

41. Kohanbash, G. et al. Isocitrate dehydrogenase mutations suppress STAT1 and CD8+ T cell accumulation in gliomas. J. Clin. Invest. 127, 1425–1437 (2017).

42. Spranger, S., Bao, R. & Gajewski, T. F. Melanoma-intrinsic β-catenin signalling prevents anti-tumour immunity. Nature 523, 231–5 (2015).

43. Chambers, A. F., Groom, A. C. & MacDonald, I. C. Dissemination and growth of cancer cells in metastatic sites. Nat. Rev. Cancer 2, 563–72 (2002).

44. Van Allen, E. M. et al. Genomic correlates of response to CTLA-4 blockade in metastatic melanoma. Science 350, 207–211 (2015).

45. Hugo, W. et al. Genomic and Transcriptomic Features of Response to Anti-PD-1 Therapy in Metastatic Melanoma. Cell 165, 35–44 (2016).

46. Tang, K.-W., Alaei-Mahabadi, B., Samuelsson, T., Lindh, M. & Larsson, E. The landscape of viral expression and host gene fusion and adaptation in human cancer. Nat. Commun. 4, 2513 (2013).

47. Angelova, M. et al. Characterization of the immunophenotypes and antigenomes of colorectal cancers reveals distinct tumor escape mechanisms and novel targets for immunotherapy. Genome Biol. 16, 64 (2015).

48. Bindea, G. et al. Spatiotemporal dynamics of intratumoral immune cells reveal the immune landscape in human cancer. Immunity 39, 782–795 (2013).

49. Rosenthal, R., McGranahan, N., Herrero, J., Taylor, B. S. & Swanton, C. deconstructSigs: delineating mutational processes in single tumors distinguishes DNA repair deficiencies and patterns of carcinoma evolution. Genome Biol. 17, 31 (2016).

50. Aran, D. et al. Widespread parainflammation in human cancer. Genome Biol. 17, 145 (2016).

